# Benchmarking single-cell foundation models for aging biology

**DOI:** 10.64898/2026.09.21.753191

**Authors:** Xiaomin Ni, Yunhao Liang, Jialin Zhu, Luoxi Zhang, Xingyu Shen, Min Yang, Ruifeng Xu, Hui Li, Rongrong Ji, Chen Jin, Shiwen Ni

**Author notes:** Correspondence (Shiwen Ni); (Chen Jin). These authors contributed equally to this work (Xiaomin Ni, Yunhao Liang, Jialin Zhu).

## Abstract

Single-cell foundation models (scFMs) provide representations of cellular states, but their utility across biological questions in aging research remains unclear. We established a benchmark of cellular representations for aging research, evaluating ten general-purpose scFMs, three aging-specific models and conventional methods across five biological questions using more than2.5 million single-cell transcriptomes. Using frozen pretrained representations, Geneformer performed best among scFMs for chronological-age prediction and age–pseudotime concordance, although 2,000 highly variable genes achieved higher mean performance. Several scFMs captured positive molecular-age shifts across three disease contexts, consistent with reported aging-associated changes. SCimilarity performed well for rare cellular state identification across out-of-distribution datasets, exceeding aging-specific models and conventional baselines. At the gene level, scGPT showed the highest recovery of reference TF–target interactions, including aging-related regulatory hubs. Overall, scFMs supported diverse aging analyses, but performance depended on the biological question, highlighting their utility for rare cellular state identification and regulatory analysis.

## Introduction

Aging is a fundamental biological process and an important contributor to declining tissue function and age-related disease susceptibility (López-Otín et al., 2023). Understanding the molecular and cellular changes that occur during aging is central to defining the mechanisms of health decline and the factors that influence disease susceptibility. Advances in single-cell transcriptomics have revealed age-associated molecular variation across cell populations and individuals, enabling the study of aging-related transcriptional programs and regulatory changes in specific cellular contexts (Emani et al., 2024; Mesecar et al., 2026; Tabula Muris Consortium, 2020). These data have enabled transcriptomic clocks to estimate age and characterize disease-associated age deviations (Mao et al., 2023; Muralidharan et al., 2025; Zhu et al., 2023), as well as approaches to identify heterogeneous senescent-cell states (Qu et al., 2025; Tao et al., 2024). Together, these approaches enable the characterization of age-associated molecular variation and heterogeneous cellular states at single-cell resolution.

Single-cell foundation models (scFMs) are pretrained on large-scale transcriptomic data and provide a new way to represent and analyze cellular and molecular variation (Cui et al., 2024; Hao et al., 2024; Theodoris et al., 2023). Their applications include cell annotation, data integration, perturbation prediction and other downstream analyses. Existing benchmarks have evaluated methodological tasks alongside biological applications, including cancer-cell identification and drug-sensitivity prediction (Wu et al., 2025). scFMs have also been used to investigate senescence- and inflammation-associated regulatory networks (Kalfon et al., 2025). However, the utility of pretrained representations across complementary aspects of aging remains insufficiently characterized. For aging, an important question is whether these representations can provide a new perspective for studying distinct aging phenotypes and whether their usefulness is consistent across biological and data contexts.

Here, we benchmark scFMs across five aging-related biological questions: chronological-age prediction, age-pseudotime concordance, disease-associated shifts in predicted age, senescent-cell identification and regulatory edge recovery. We compare ten general-purpose scFMs with one aging-specific model, two senescence-specific models and conventional baselines, using datasets comprising more than 2.5 million single-cell transcriptomes. The pretrained models are kept frozen, and supervised predictors are trained on their representations. This design allows us to examine the usefulness of pretrained representations across distinct aging phenotypes, including where scFMs provide useful information, where conventional features remain effective, and how their performance varies across models and biological questions.

## Results

### A benchmark framework for evaluating scFMs in aging research

We established a five-task benchmark to examine which aspects of aging biology are accessible from pretrained scFM representations. The tasks assessed donor-level age prediction, age-pseudotime concordance, disease-associated age shifts, senescent-cell identification and regulatory edge recovery (Fig. 1A). Cell representations were used for the first four tasks and gene representations for the regulatory analysis. Foundation models were kept frozen; age regressors and senescence classifiers were trained on the extracted representations. Conventional and specialized methods provided comparators for each biological question.

**Figure 1.**
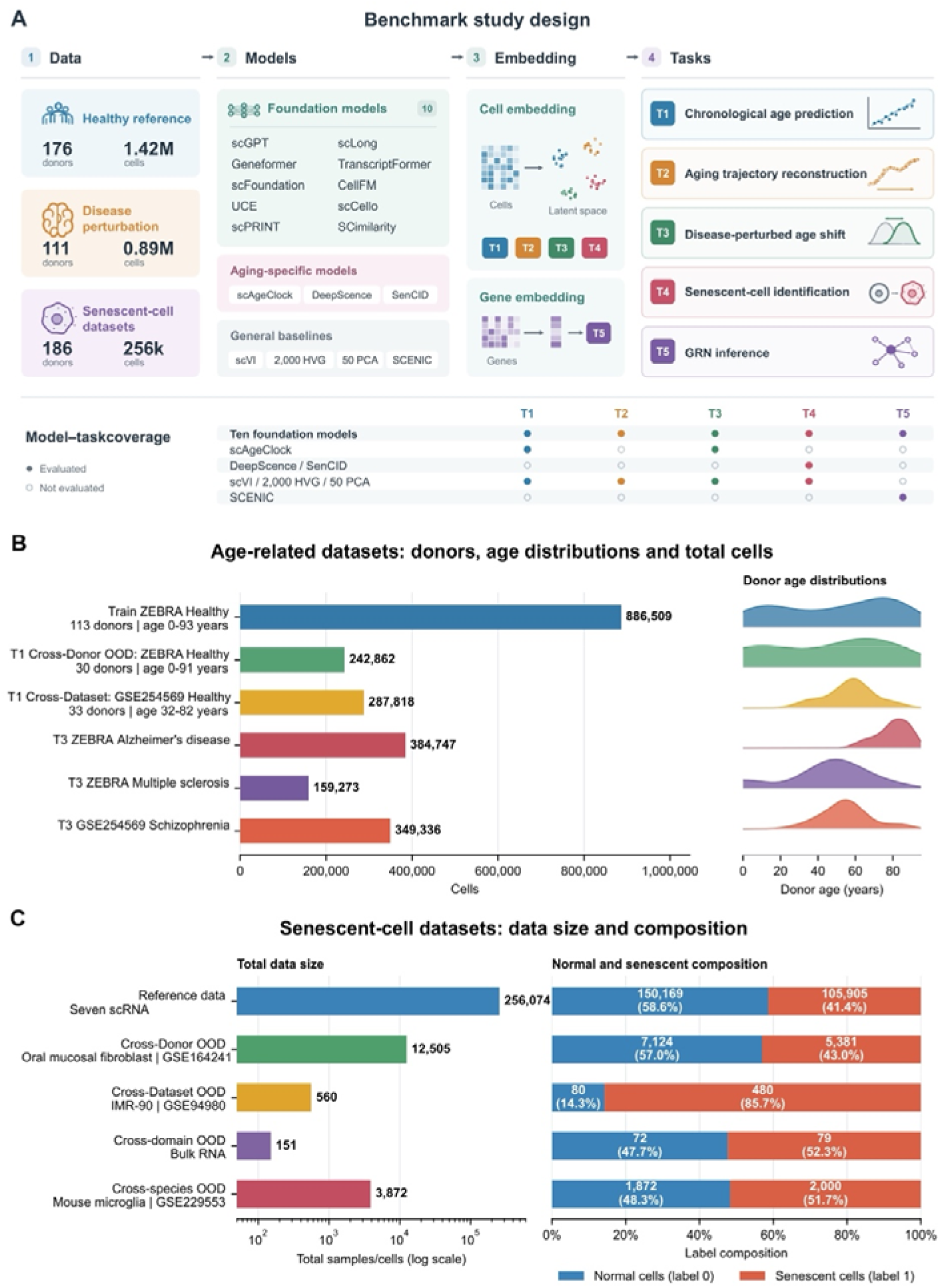
Benchmark design and dataset composition for evaluating single-cell foundation models in aging. (A) Benchmark datasets, models, embedding strategies, and five downstream tasks. Filled circles denote evaluated model–task combinations. (B) Cell numbers and donor-age distributions across healthy and disease-perturbed datasets in Tasks 1, 2, 3, and 5. (C) Task 4 reference collection (256,074 cells) and independent evaluation datasets; the classifier development subset contained 11,280 cells. AD, Alzheimer’s disease; MS, multiple sclerosis; GRN, gene regulatory network.

The benchmark assembled datasets comprising more than 2.5 million single-cell transcriptomes. We evaluated ten general-purpose scFMs (scGPT, Geneformer, scFoundation, UCE, scPRINT, scLong, TranscriptFormer, CellFM, scCello and SCimilarity), three specialized models (scAgeClock, DeepScence and SenCID; Xie, 2026; Qu et al., 2025; Tao et al., 2024) and three conventional baselines (scVI, 2,000 highly variable genes [HVGs] and 50 principal components [PCs]). SCENIC served as the baseline for regulatory edge recovery. Age predictors were developed using 113 healthy ZEBRA donors and evaluated on held-out donors, independent GSE254569 controls and three disease cohorts (Fig. 1B). Age-pseudotime concordance was assessed across six healthy ZEBRA cell types (Supplementary Fig. S1A; Supplementary Table S7). The senescence reference collection contained 256,074 cells from seven datasets; classifiers were developed using a selected set of 11,280 cells and evaluated across donors, datasets, measurement domains and species (Fig. 1C). Dataset composition and model coverage are summarized in Supplementary Fig. S1 and Supplementary Tables S1 and S2.

### scFM representations retain age information but do not consistently outperform conventional features

We first assessed whether pretrained scFM representations retained information useful for chronological-age prediction. At the cell level, PCCs across the ten scFMs ranged from 0.647 to 0.919 in held-out healthy ZEBRA samples, decreasing to 0.080–0.391 in the independent GSE254569 cohort. Corresponding MAEs increased from 6.841–16.665 to 26.363–35.542 years. After aggregating predictions by donor, PCCs ranged from 0.722 to 0.943 in ZEBRA and from 0.467 to 0.718 in GSE254569. Averaged across these two cohorts, Geneformer achieved the highest donor-level PCC among scFMs (0.831), whereas the 2,000-HVG baseline yielded a higher mean PCC of 0.848. The aging-specific model scAgeClock had a mean donor-level PCC of 0.232, indicating limited predictive performance in the evaluated brain cohorts. Thus, pretrained scFMs retained age-associated information, but conventional expression features provided stronger overall prediction, and aging-specific model development did not ensure effective transfer to these cohorts (Fig. 2A,B; Supplementary Tables S3 and S4).

**Figure 2.**
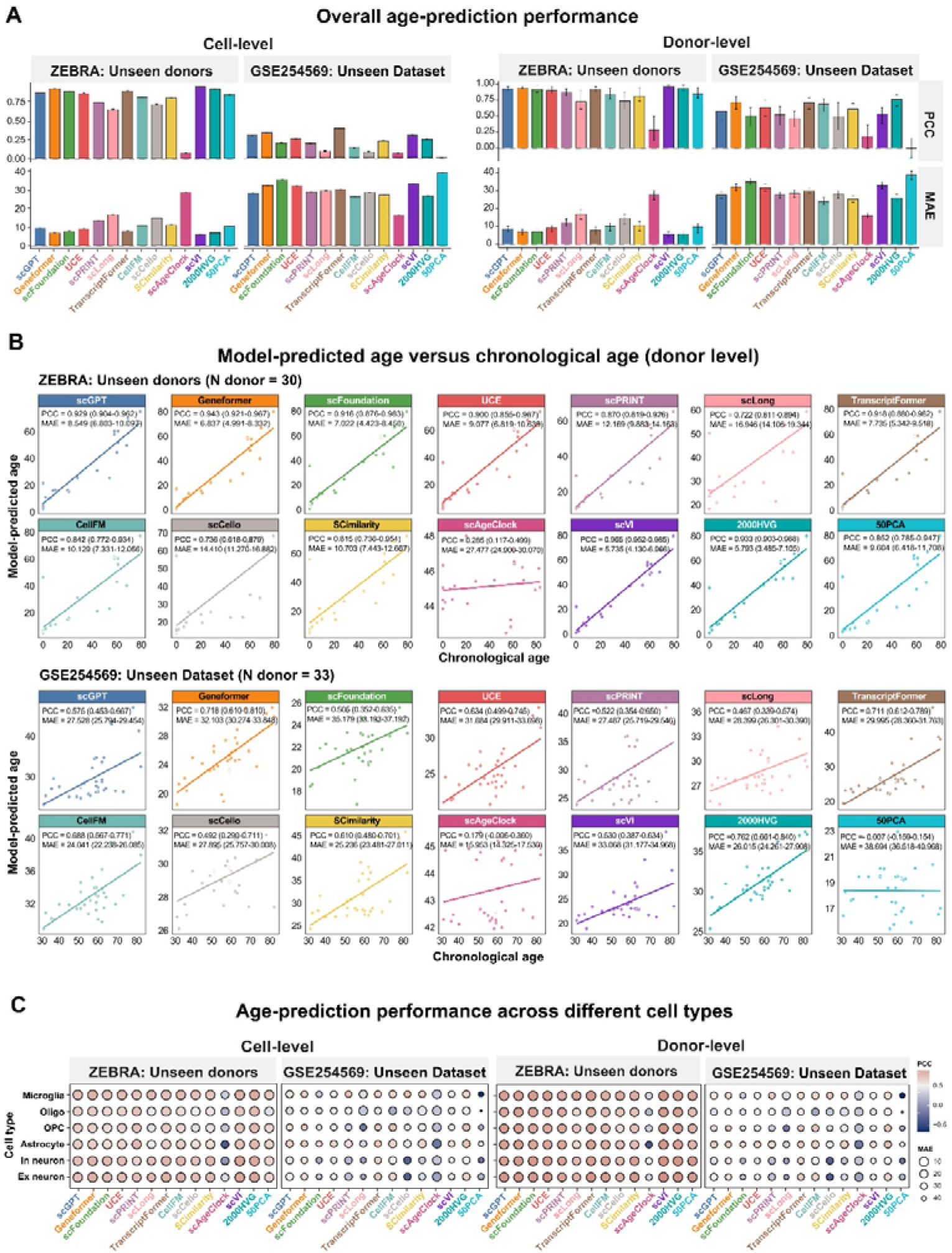
Benchmarking single-cell foundation models for age prediction. (A) Overall age-prediction performance on unseen donors and unseen datasets, evaluated at both the cell level (left) and donor level (right). Performance is assessed using Pearson correlation coefficient (PCC) and mean absolute error (MAE). (B) Scatterplots of model-predicted age versus chronological age at the donor level for unseen donors and unseen datasets. Mean PCC and MAE with 95% confidence intervals (CIs) are shown in the upper-left corner of each panel. (C) Age-prediction performance across different cell types in unseen donors and unseen datasets, evaluated at the cell level (left) and donor level (right) using PCC and MAE.

Cell-type-stratified analyses further showed that age-prediction performance varied across neuronal and glial populations (Fig. 2C). For Geneformer, donor-level PCCs in healthy ZEBRA samples were 0.970 in excitatory neurons and 0.921 in microglia (Supplementary Fig. S2), decreasing to 0.594 and 0.645, respectively, in GSE254569. Thus, the reduction in predictive performance across cohorts was also observed within individual brain cell populations (Supplementary Tables S5 and S6).

### Age–pseudotime concordance is cell-type dependent and does not consistently exceed HVG features

We next asked whether model-derived representations could recover continuous age-related cellular trajectories. For each cell type, we constructed pseudotemporal trajectories and quantified their concordance with chronological age using Pearson correlation. The ability to recover age-ordered trajectories varied substantially across representations and cellular identities. Notably, the conventional 2,000-HVG representation showed the strongest overall concordance with chronological age (mean Pearson r = 0.736), exceeding all foundation models, whose mean correlations ranged from 0.284 to 0.595. Geneformer performed best among the foundation models (mean r = 0.595), whereas scPRINT showed the weakest association (r = 0.284) (Fig. 3A,B; Supplementary Table S7).

**Figure 3.**
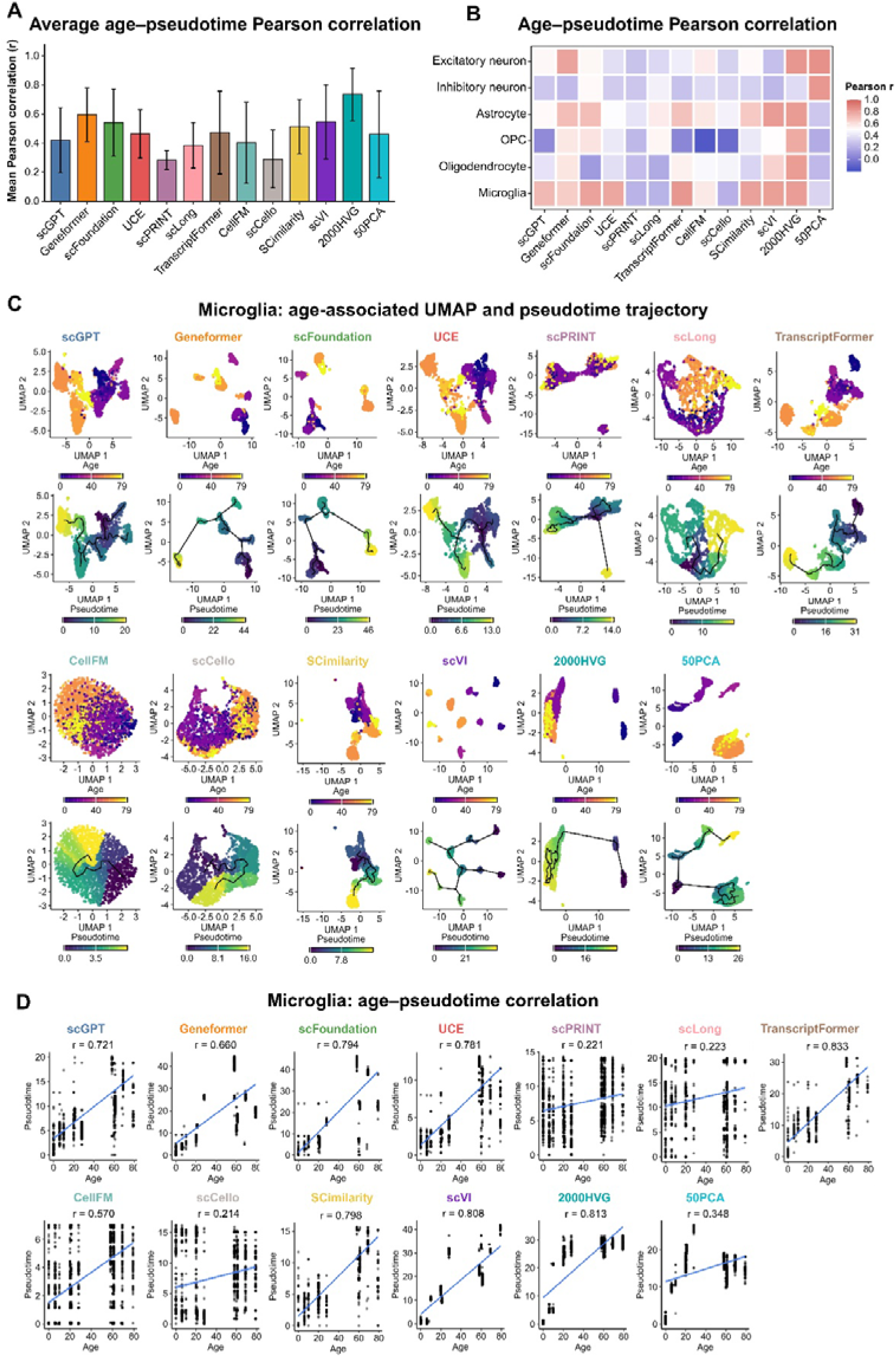
Benchmarking age-associated pseudotime representations across single-cell models. (A) Average age–pseudotime Pearson correlation across cell types. (B) Age–pseudotime Pearson correlation across cell types and models. (C) Age-associated UMAP and pseudotime trajectory of Microglia. (D) Age–pseudotime relationship in Microglia.

The relative performance of representations also varied across cell types. Geneformer showed a strong age–pseudotime association in excitatory neurons (r = 0.792), while scFoundation and TranscriptFormer performed particularly well in microglia (r = 0.794 and 0.833, respectively); SCimilarity also showed a strong association in microglia (r = 0.798). In contrast, several representations showed weak age–pseudotime concordance in oligodendrocytes and oligodendrocyte precursor cells (OPCs). Importantly, 2,000 HVGs remained competitive across diverse cell types and frequently outperformed foundation-model representations, indicating that no foundation model consistently outperformed conventional transcriptomic representations across cellular contexts (Fig. 3B-D; Supplementary Fig. S3; Supplementary Table S7).

### scFM representations show disease-dependent changes in age-associated signals

To examine disease-associated changes in the age-related signal, we applied the age predictors to Alzheimer’s disease (AD), multiple sclerosis (MS) and schizophrenia cohorts. Cell-level PCCs among scFMs were 0.029–0.370 in AD and −0.299–0.381 in MS, below their values in healthy ZEBRA evaluation. In schizophrenia, PCCs of 0.432–0.602 were higher than in GSE254569 controls. ΔMAE was positive for most methods in AD and MS and negative for all methods in schizophrenia. Disease was therefore associated with different changes in prediction accuracy across cohorts (Fig. 4A–C; Supplementary Tables S3 and S8).

**Figure 4.**
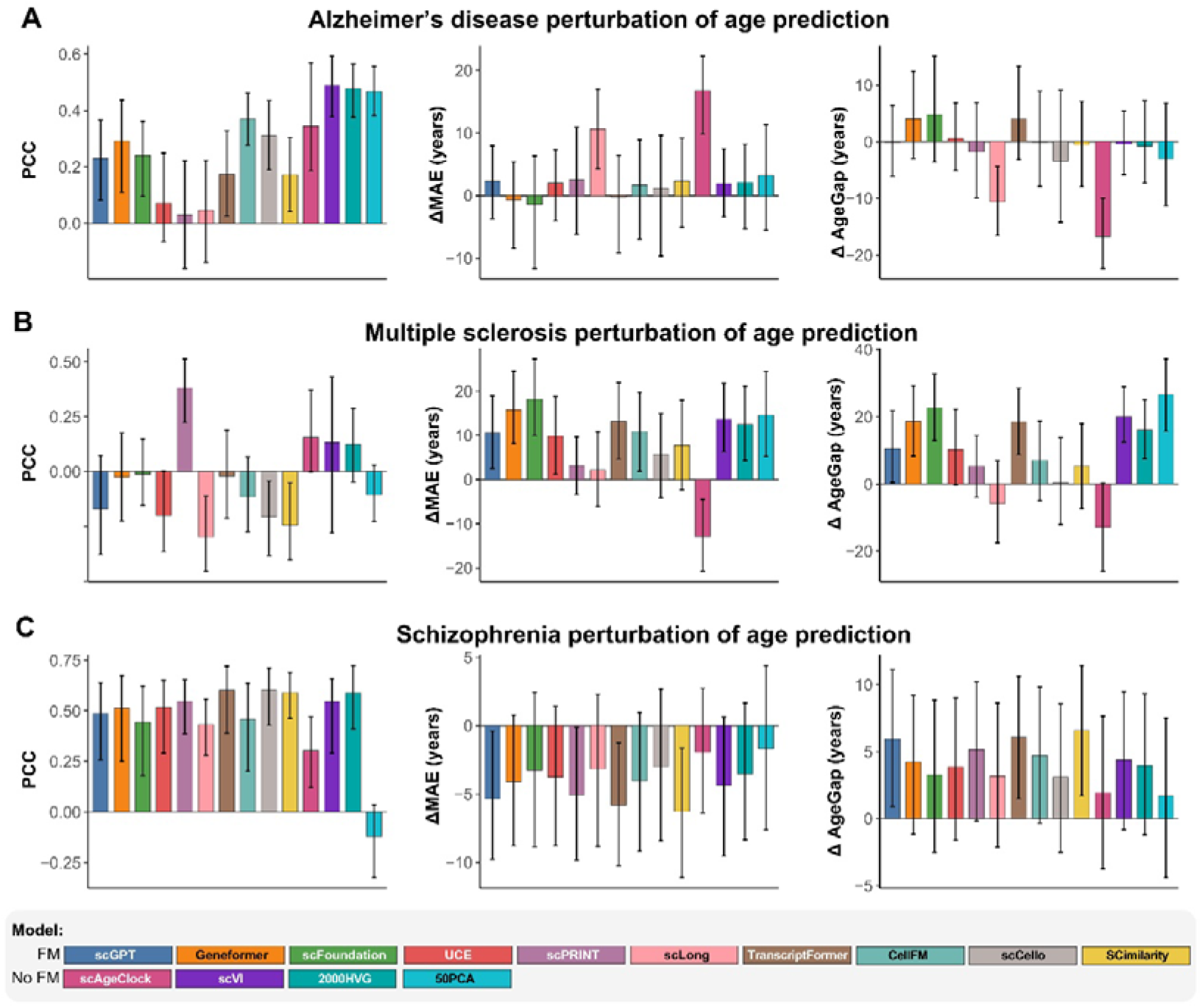
**Disease-associated shifts in transcriptomic age prediction**. (A–C) Alzheimer’s disease, multiple sclerosis and schizophrenia, respectively. Within each row, panels show PCC between predicted and chronological age, ΔMAE and ΔAgeGap from left to right. Differences are calculated as disease minus dataset-matched control values; ΔAgeGap compares mean donor-level age gaps. Bars show point estimates and error bars indicate 95% confidence intervals.

We separately examined the disease–control difference in mean AgeGap (ΔAgeGap, with AgeGap defined as predicted minus chronological age). In AD, four scFMs had positive ΔAgeGap: Geneformer (+4.06 years), scFoundation (+4.78), UCE (+0.60) and TranscriptFormer (+4.00). Positive differences occurred for 12 of 14 methods in MS and all 14 in schizophrenia. In schizophrenia, the relative shift accompanied reduced prediction error against a background of age underestimation in controls (Fig. 4A–C; Supplementary Tables S3 and S8).

Across the three diseases, four methods showed positive mean differences in all contexts, eight in two contexts and two only in schizophrenia. These patterns distinguish the direction of a disease-associated age shift from the accuracy of chronological-age prediction. Cell-type-specific estimates are reported in Supplementary Fig. S4 and Supplementary Table S9, with directional summaries in Supplementary Table S18.

### scFMs support identification of rare and heterogeneous senescent-cell states across out-of-distribution settings

We then assessed whether cell representations supported recognition of senescence-associated states across datasets. UMAPs revealed substantial differences in the separation of senescent and non-senescent cells across model representations, with some models showing clear state separation while others showed extensive overlap. These differences were reflected in classification performance. On unseen donors, foundation models achieved F1 scores of 0.401–0.820, with SCimilarity performing best (F1 = 0.820, sensitivity = 0.713, PPV = 0.965). UCE, TranscriptFormer and scCello also showed strong performance (F1 = 0.790–0.804), generally exceeding conventional baselines and most senescence-specific models (Fig. 5A,B; Supplementary Table S10).

**Figure 5.**
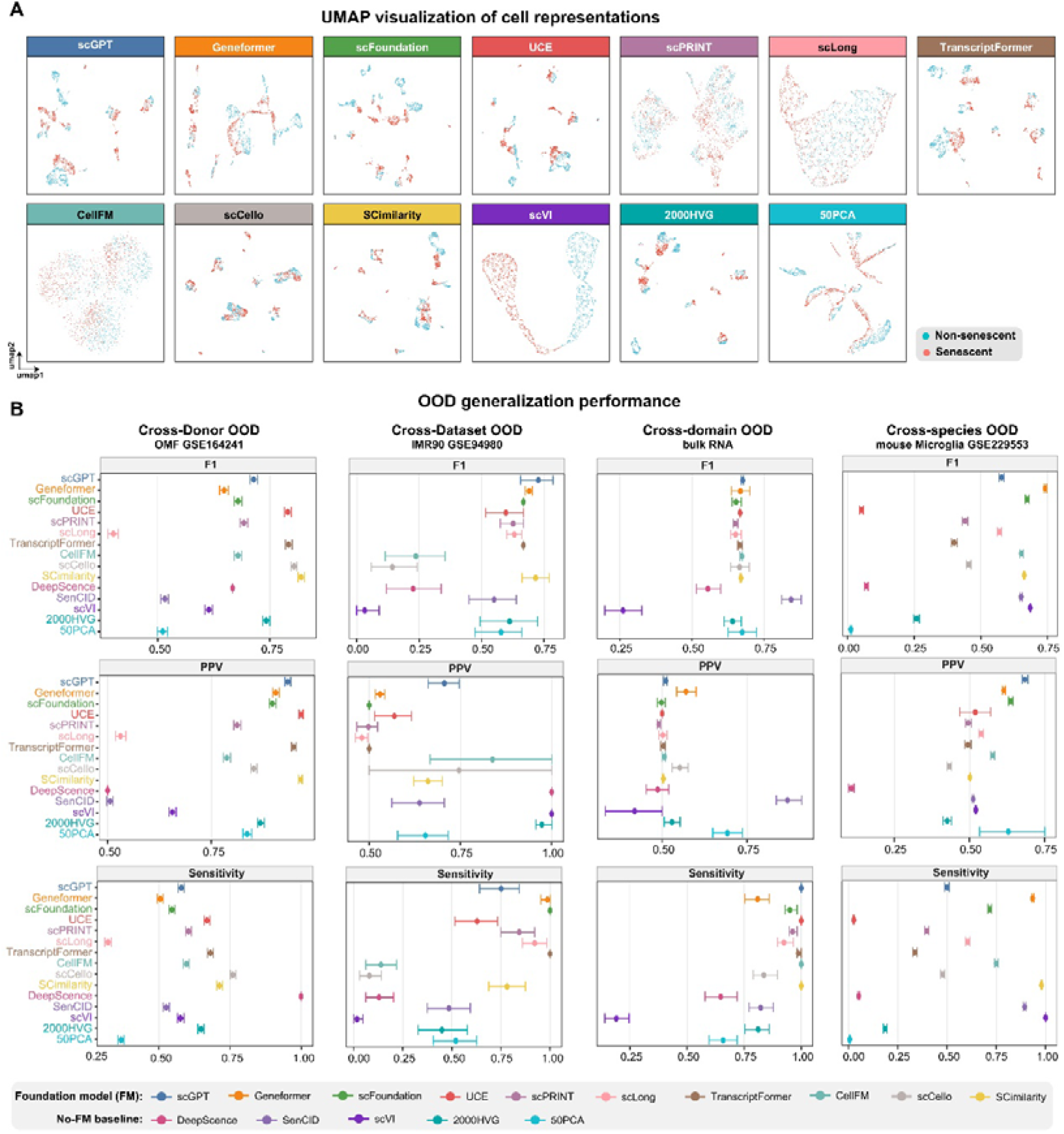
**Generalization of single-cell foundation models for senescent-cell identification across multiple out-of-distribution settings**. (A) UMAP visualization of cell representations under the cross-donor OOD setting. (B) Senescent-cell identification performance across four OOD settings: cross-donor, cross-dataset, cross-domain using bulk RNA-seq data, and cross-species. Performance is evaluated using F1 score, positive predictive value (PPV), and sensitivity. Additional classification metrics, including PR-AUC, ROC-AUC, and Cohen’s kappa, are provided in Supplementary Fig. S5. Comprehensive model- and dataset-level results are provided in Supplementary Table S10.

Generalization to unseen datasets was more heterogeneous, with F1 scores ranging from 0.143–0.727. Geneformer, scGPT and SCimilarity retained relatively strong performance (F1 = 0.690–0.727), although sensitivity and PPV showed substantial trade-offs across models. In bulk RNA-seq, most foundation models showed high sensitivity but lower PPV, whereas the bulk-oriented SenCID achieved the highest overall performance (F1 = 0.844, sensitivity = 0.822, PPV = 0.869), highlighting the challenge of transferring senescence representations from single-cell to bulk data. Cross-species transfer to mouse further accentuated model differences, with F1 scores ranging from 0.052–0.741; Geneformer performed best (F1 = 0.741, sensitivity = 0.934, PPV = 0.614), whereas UCE showed a marked decline (Fig. 5B; Supplementary Fig. S5; Supplementary Table S10).

Overall, senescent cell identification showed substantial heterogeneity across foundation models and distribution shifts. Notably, SCimilarity, scGPT and Geneformer showed strong performance across multiple OOD settings and, in several scenarios, outperformed both senescence-specific models and conventional baselines. However, no single foundation model consistently dominated across all distribution shifts, indicating that model performance depended strongly on the type of distribution shift.

### scFM gene representations recover reference regulatory relationships and reveal recurrent network hubs

We next assessed whether gene-level representations extracted from scFMs could recover reference TF–target interactions. Pre-motif AUPRC was evaluated on a common universe of 163,280 candidate TF–target pairs. scGPT achieved the highest AUPRC (0.006198), followed by SCimilarity (0.005358), scLong (0.004325) and UCE (0.004134). These four representations had higher point estimates than the SCENIC baseline at the GRNBoost2 stage (0.003481), whereas the remaining six scFMs yielded lower AUPRCs (Fig. 6A; Supplementary Fig. S7; Supplementary Table S17).

**Figure 6.**
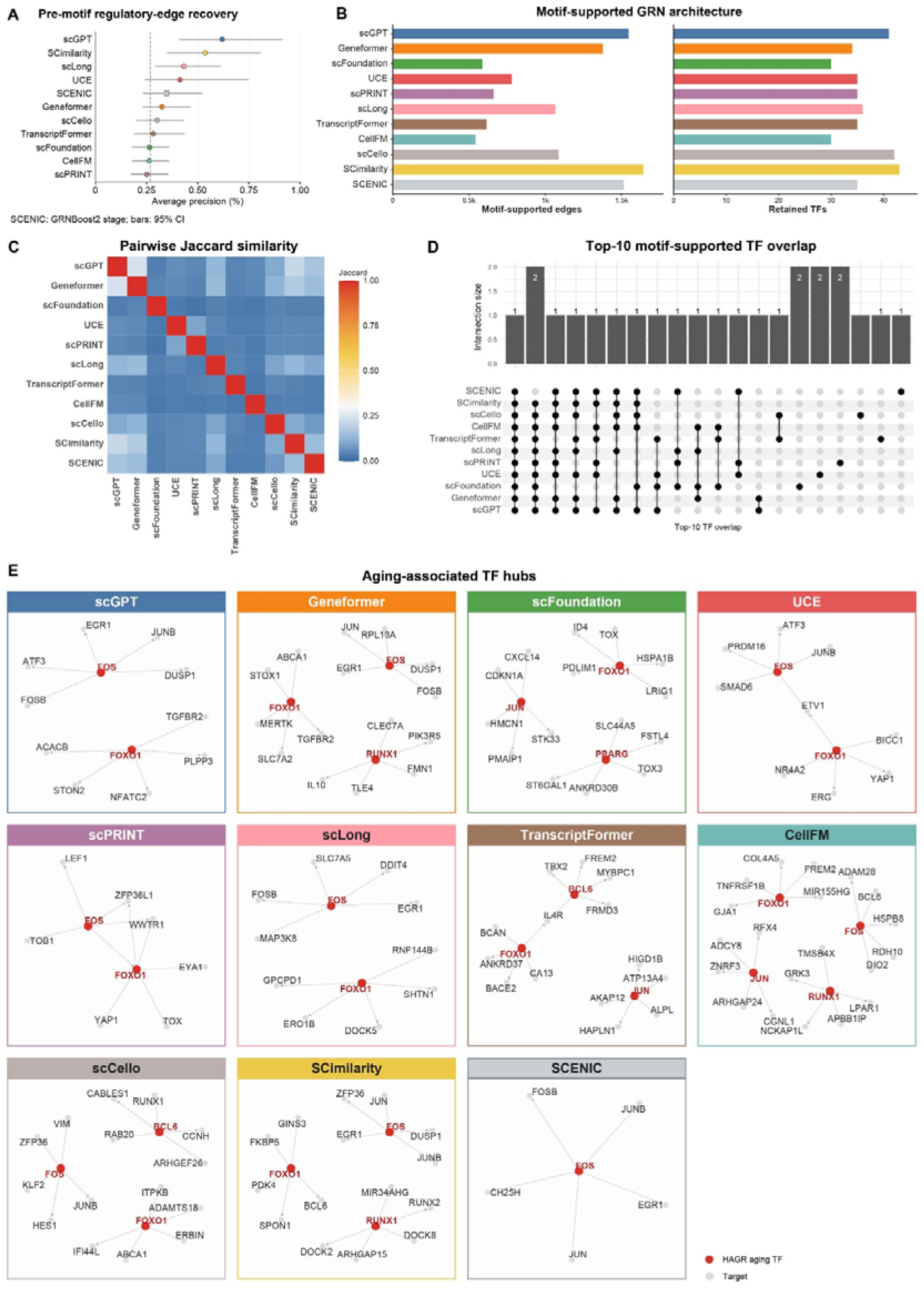
Cross-model comparison of transcriptional regulatory architectures and aging-associated regulatory programs. (A) Regulatory-edge recovery across scFMs and SCENIC, evaluated by pre-motif AUPRC. (B) Numbers of motif-supported edges and retained TFs; SCENIC is shown in light gray. (C) Pairwise Jaccard similarity of complete motif-supported edge sets. (D) UpSet plot of 20 membership patterns among the top 10 TF hubs across networks. (E) TFs recovered among each model’s top 10 hubs and their five highest-scoring motif-supported targets.

An analysis of motif-supported network structure yielded 542–1,643 retained edges and 30–43 TFs per scFM (Fig. 6B). Edge-set Jaccard similarities between scFM-derived networks and matched-input SCENIC ranged from 0.033 to 0.162, indicating limited overlap in inferred interactions (Fig. 6C). Despite these edge-level differences, pairs of scFM networks shared 4–8 of their top ten TF hubs (Fig. 6D; Supplementary Table S11). Recurrent hubs included FLI1, CEBPD, IKZF1, IRF8, FOXO1 and FOS. Each scFM network contained 2–4 known aging-associated TFs among its top ten hubs; CellFM’s four annotated hubs were FOS, FOXO1, RUNX1 and JUN (Fig. 6E). The all-expressed-gene SCENIC sensitivity analysis is reported in Supplementary Fig. S6 and Supplementary Table S12.

### Cross-task comparisons reveal where scFMs are useful in aging research

Across the five tasks, no single scFM performed consistently best (Fig. 7). The 2,000-HVG baseline ranked first among all eligible methods for donor-level age prediction and age–pseudotime concordance, with mean PCCs of 0.848 and 0.736, respectively. Geneformer achieved the highest mean PCCs among scFMs (0.831 and 0.595). Its Transformer encoder uses expression-ranked gene tokens and was pretrained with masked gene prediction (Supplementary Table S13). For disease-associated age shifts, Geneformer, scFoundation, UCE and TranscriptFormer showed positive ΔAgeGap estimates across all three disease contexts, in the direction of previously reported aging-related changes in these disorders, and shared the highest T3 rank for directional consistency.

**Figure 7.**
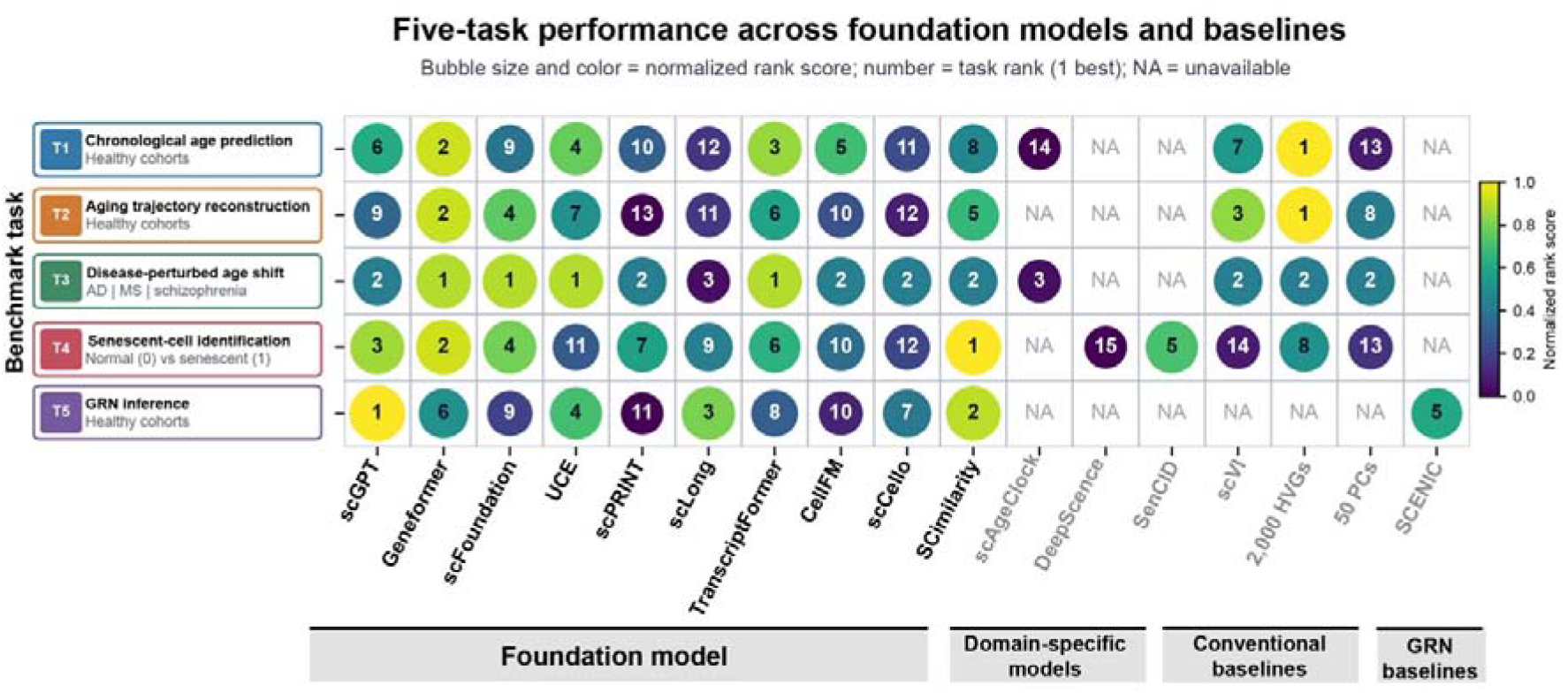
Cross-task performance profiles of foundation models and baselines. Bubble size and color indicate normalized average-rank scores. Numbers show dense task ranks (1 = best; ties are numbered consecutively, for example, 1, 1, 2); NA denotes unavailable results.

SCimilarity achieved the highest mean F1 for senescent-cell identification across the four settings (0.717), exceeding conventional baselines (0.398–0.564) and senescence-specific models (0.380–0.640). Its representations were generated by an MLP-based model trained with reconstruction and triplet metric-learning objectives (Supplementary Table S13). For GRN edge recovery, scGPT’s learned gene-token embeddings achieved the highest reported AUPRC (0.006198), followed by SCimilarity (0.005358), scLong (0.004325) and UCE (0.004134); SCENIC ranked fifth (0.003481). Geneformer had the highest five-task mean normalized rank score (0.856), followed by SCimilarity (0.689), TranscriptFormer (0.656) and scGPT (0.622).

These performance profiles spanned models with different architectures, expression encodings and pretraining strategies (Supplementary Table S13), whose individual effects were not isolated in this benchmark. Overall, pretrained representations showed phenotype-dependent advantages, while conventional HVG features yielded stronger aggregate performance in chronological-age prediction and age–pseudotime concordance.

## Discussion

Aging provides an important context for testing whether single-cell foundation models (scFMs) can support real biological questions, because aging is reflected in continuous, heterogeneous and multidimensional changes at the molecular and cellular levels. Existing benchmarks have mainly evaluated models on specific methodological tasks. Our benchmark instead examines pretrained representations in the context of aging biology and asks whether they can support different aging-related phenotypes. Without aging-specific fine-tuning, the representations supported analyses of chronological age, disease-associated molecular changes and senescence-associated cellular states, but their performance was not consistent across models or questions. Conventional expression features were also more effective in some analyses. Taken together, these findings suggest that the biological information provided by scFMs is not a single and uniform property, but differs across the aging phenotype being studied.

Senescent-cell identification is one important example. Senescent cells show diverse transcriptional and secretory phenotypes across cell types and physiological contexts, and they can affect neighboring cells through paracrine signaling (Gorgoulis et al., 2019; Wang et al., 2024). When senescent cells are rare in tissues, their contribution to tissue phenotypes can be difficult to recognize from their abundance alone. Identifying candidate senescent states may therefore help determine which cellular phenotypes are associated with tissue dysfunction and which are responsive to senescence-targeting interventions. The present findings suggest that scFM representations may be useful for this purpose, particularly for heterogeneous and relatively rare cellular states. Their biological interpretation, however, still needs to be supported by independent phenotypic evidence.

At the gene level, scFM representations provide another way to examine regulatory programs related to aging. Gene co-expression and motif evidence can support the interpretation of candidate TF–target relationships (Aibar et al., 2017). Their relevance to the cellular state of interest could be further assessed using tissue-specific expression and chromatin accessibility. These predicted relationships should be viewed as hypotheses rather than direct evidence of regulatory mechanisms. Perturbation experiments in young and aged cells could then test whether candidate regulators affect cellular functions or responses to intervention, and whether such effects depend on age. In this way, gene representations may help move from age-associated transcriptional changes to candidate regulatory mechanisms that can be tested experimentally.

The conclusions of this study are based on frozen pretrained representations and the downstream analyses evaluated here. The foundation models were not fine-tuned for the individual tasks, and our results therefore reflect the information available from their pretrained representations under the conditions examined here. Task-specific fine-tuning may further improve their performance for particular aging-related questions and represents an important direction for future application. The age-related analyses focused on brain cohorts, while gene-representation extraction, senescence annotations and the choice of reference regulatory networks also affect the scope of the findings. Within these conditions, our benchmark shows that scFMs can support some questions in aging research, while also revealing differences in the biological information they provide.

Senescent-cell identification and regulatory hypothesis generation are two areas where these representations may help link age-associated molecular changes to cellular states and regulatory processes.

## Methods

### Dataset collection and processing

#### Data sources and eligibility criteria

Publicly available single-cell RNA-seq datasets relevant to the analyses were collected from the original studies. We included datasets with processed gene-count matrices and the corresponding donor- and cell-level annotations required for each analysis. We used the source studies’ quality-control criteria. Dataset accessions, sample counts and references are listed in Supplementary Table S1.

#### Healthy cohorts for age-related analyses

Age-related analyses used ZEBRA cortex and non-cortex samples (Flotho et al., 2024) and an independent orbitofrontal-cortex cohort, GSE254569 (Fröhlich et al., 2024). Model development included 113 healthy ZEBRA donors aged 0–93 years (886,509 original cells; 139,850 benchmark cells). Healthy evaluation included 30 held-out ZEBRA donors aged 0–91 years (242,862 original cells; 29,127 benchmark cells) and 33 GSE254569 control donors aged 32–82 years (287,818 original cells; 19,313 benchmark cells).

#### Disease cohorts for age-shift analysis

Disease evaluation included 45 ZEBRA donors with Alzheimer’s disease (23,874 benchmark cells), 30 with multiple sclerosis (32,297 cells), and 36 GSE254569 donors with schizophrenia (20,878 cells). Controls were selected from the corresponding source cohort and were not pooled across datasets.

#### Gold-standard datasets for senescent-cell classification

Seven published datasets comprising 256,074 cells were assembled as a reference collection for senescent-cell classification: HCA2 fibroblasts, GSE119807 (Tang et al., 2019); IMR-90 fibroblasts, GSE115301 (Teo et al., 2019); HUVECs, GSE102090 (Zirkel et al., 2018); pulmonary epithelial cells, GSE190889 (Heinzelmann et al., 2022); oral mucosal fibroblasts, GSE164241 (Williams et al., 2021); and WI-38 fibroblasts, GSE226225 (Wechter et al., 2023). The classifier development set contained 11,280 cells: 5,607 non-senescent and 5,673 senescent cells, encoded as 0 and 1. Classifiers were tested on held-out GSE164241 donors (12,505 cells), independent IMR-90 data from GSE94980 (560 cells; Aarts et al., 2017), 151 bulk RNA-seq samples from 17 studies, and mouse microglia from GSE229553 (3,872 cells; Ng et al., 2023). Independent-test labels were used only for evaluation.

#### Metadata harmonization and cell sampling

Brain cell annotations were mapped to six cell types: excitatory neurons, inhibitory neurons, astrocytes, oligodendrocyte precursor cells, oligodendrocytes and microglia. Other cell types were included only in analyses of the full dataset. Donors were assigned to development or independent-test cohorts before sampling, with up to 2,000 cells retained per donor and cell type. The internal training–validation split was performed at the cell level, as described below.

#### Extraction of pretrained cell representations

Gene identifiers were matched to each model’s vocabulary, and expression matrices were processed using the corresponding inference pipeline. Pretrained encoder weights were held fixed while downstream predictors were trained. Embeddings were linked to sample metadata by cell identifier. Checkpoints, normalization, tokenization, pooling and embedding dimensions are listed in Supplementary Table S14.

Cell representations were extracted from scGPT (Cui et al., 2024), Geneformer (Theodoris et al., 2023), scFoundation (Hao et al., 2024), UCE (Rosen et al., 2026), scPRINT (Kalfon et al., 2025), scLong (Bai et al., 2026), TranscriptFormer (Pearce et al., 2026), CellFM (Zeng et al., 2025), scCello (Yuan et al., 2024) and SCimilarity (Heimberg et al., 2025), yielding one fixed-dimensional vector per cell.

CellFM inputs were normalized by library-size scaling and rounding before sparse storage. Up to 2,048 expressed genes were retained per cell; when this limit was exceeded, genes were sampled using probabilities derived from log-transformed expression. The frozen CellFM 80M encoder provided a 1,536-dimensional CLS representation. scPRINT used medium v1.5 with up to 2,000 expression-informed gene tokens and 256-dimensional outputs; mouse-derived inputs were mapped to human orthologues while retaining their source-species annotation.

#### Extraction of gene representations

Gene representations were extracted from pretrained models or their associated resources without further training. Depending on the model, these comprised gene-token embeddings, positional embeddings, protein-derived vectors, Gene2Vec vectors, decoder coefficients or contextual gene embeddings. Gene identifiers were harmonized across models; extraction and mapping procedures are listed in Supplementary Table S14. Regulatory-edge recovery was assessed using the shared gene set defined below.

#### Conventional expression-based representations

##### 2,000 highly variable genes

The 2,000 highly variable genes (HVGs) were selected from the training data using variance-stabilizing transformation (Stuart et al., 2019) after standard normalization. Evaluation cells were normalized using the same procedure and aligned to the training-derived HVG set, with absent genes zero-filled. The resulting 2,000-dimensional vectors were used as the HVG baseline.

##### 50 principal components

PCA (Jolliffe and Cadima, 2016) was fitted to the normalized training data restricted to the 2,000 training-derived HVGs, and the first 50 components were retained. Evaluation cells were projected using the training-derived scaling parameters and PCA rotation without refitting. The resulting 50-dimensional vectors were used as the PCA baseline.

##### scVI

scVI (Lopez et al., 2018) was fitted separately to each input cohort using raw counts and 30 latent dimensions, with scvi-tools defaults unless otherwise specified (Gayoso et al., 2022). Because evaluation cohorts contributed to representation fitting, scVI was treated as a transductive baseline.

#### Downstream prediction framework

##### Model architecture and optimization

Multilayer perceptrons were trained on frozen cell embeddings or conventional expression features. Age predictors had hidden layers of 512 and 128 units, except for TranscriptFormer (512, 256 and 128) and conventional baselines (256 and 128). Hidden layers used normalization, ReLU activation and dropout, followed by a single output unit. Features were standardized using means and standard deviations from the development cohort; these values were also used for validation and independent-test data.

Predictors were optimized with AdamW (Loshchilov and Hutter, 2019), using mean squared error for age regression and binary cross-entropy with logits for senescence classification. Model-specific training settings were used, and early stopping retained the checkpoint with the lowest validation loss. Senescence probabilities were obtained by sigmoid transformation and binarized at 0.5, without tuning on independent-test labels.

##### Internal validation and independent evaluation

Development cells were divided into training and validation sets at a 9:1 ratio, with stratification by senescence label for classification. Internal validation was used to select the checkpoint and was not an independent-donor test. Trained predictors were then applied to independent cohorts without retraining.

#### Task 1: chronological-age prediction

Cells without valid age annotations were excluded. Development-data partitions were pooled to train one age predictor per representation, without fitting separate models for individual partitions or cell types. Predictors trained on healthy ZEBRA cells were applied to held-out healthy ZEBRA and GSE254569 control cells. Predictions were subsequently analysed by cell type, sex, tissue and donor. Donor-level predictions were calculated by averaging predicted ages across each donor’s cells, either across all cells or separately within each of the six shared cell types. scAgeClock was included as an aging-specific comparator (Xie, 2026).

Age prediction was assessed using Pearson correlation coefficient (PCC), Spearman correlation, mean absolute error (MAE), root mean squared error, R², concordance correlation coefficient, intraclass correlation coefficient (ICC), median absolute error and age gap. ICC used a two-way, absolute-agreement, single-measure definition. Age gap was predicted minus chronological age.

#### Task 2: cell-type-specific age trajectories

Trajectories were inferred separately for each cell type in the healthy ZEBRA training cohort. Cells without age or cell-type annotations and cell types with fewer than 1,000 cells were excluded. Neighborhood graphs and UMAP coordinates were computed from each representation (minimum distance, 0.01; McInnes et al., 2018). Coordinates were matched by cell identifier before trajectory inference with Monocle 3 (Cao et al., 2019); preprocessing retained up to 50 principal components.

Root cells were selected from cells with the minimum chronological age, and pseudotime was calculated as graph distance from the root. Chronological age was thus used to orient the trajectory. The primary measure was signed PCC between chronological age and pseudotime among cells with finite pseudotime values; Spearman correlation and linear-model R² were secondary measures.

#### Task 3: disease-associated shifts in predicted age

Age predictors trained on healthy ZEBRA donors were applied without retraining to Alzheimer’s disease, multiple sclerosis and schizophrenia cohorts. PCC between predicted and chronological age was calculated within each disease cohort. Comparisons used controls from the same dataset and included only donors within the age range shared by the disease and control groups.

Age gap was defined as predicted minus chronological age. ΔMAE was defined as MAE(disease) − MAE(control). Mean age gap was first calculated within donor, and ΔAgeGap was then defined as the difference between the mean donor-level age gap in the disease and control groups. Positive ΔAgeGap indicated older predicted age relative to chronological age in disease. Previous reports of transcriptomic age acceleration in Alzheimer’s disease and schizophrenia (Muralidharan et al., 2025) and glial epigenetic age acceleration in progressive multiple sclerosis (Kular et al., 2022) motivated the expected positive direction. For cross-task ranking, T3 was the equally weighted fraction of the three disease contexts with ΔAgeGap > 0. Zero or negative estimates contributed 0; missing estimates remained missing. Effect magnitudes, 95% confidence intervals and the number of intervals entirely above zero were reported separately (Supplementary Table S18) and did not break ties. This endpoint measured directional agreement, not statistically established biological age acceleration. Mean signed ΔAgeGap was retained as a sensitivity endpoint.

#### Task 4: senescent-cell identification

For each scFM, a binary classifier was trained on the 11,280-cell development set while encoder weights remained fixed. Classifiers used the same hidden layers as the corresponding age predictors, with a binary logit replacing the regression output. Standardization parameters, classifier weights and decision thresholds were retained when testing held-out donors, independent datasets, bulk RNA-seq and mouse microglia. Bulk RNA-seq predictions were evaluated at the sample level.

DeepScence (Qu et al., 2025) was fitted separately to each dataset after removing genes with no detected counts and denoising expression with Deep Count Autoencoder. Human or mouse CoreScence gene sets were used as appropriate, with mouse gene-symbol capitalization corrected where needed. Senescence labels, cell-type annotations and batch labels were not supplied during fitting. Scores were converted to binary calls using the official binarization procedure, without adjusting thresholds for individual datasets.

For SenCID (Tao et al., 2024), gene names were made unique and supplied gene-symbol columns were used when available. Released SID1–SID6 and recommended-SID models generated continuous scores and binary calls using the 0.5 threshold. DCA denoising was disabled by default. Ground-truth labels were retained only for evaluation.

Performance was assessed separately in each test setting using F1, positive predictive value, sensitivity, specificity, Cohen’s kappa, AUROC and AUPRC. F1 was used for cross-task comparison. Accuracy, balanced accuracy, Matthews correlation coefficient and confusion-matrix counts were reported where available. Metrics requiring both classes were not used to compare performance in datasets containing only one class.

#### Task 5: regulatory-edge recovery and network annotation

##### Reference-edge recovery

Regulatory-edge recovery was assessed on gene embeddings built from the 2000 HVGs (Wu et al., 2025), using 1,571 genes and 104 TFs shared across the ten scFM gene sets, which yielded 163,280 directed TF–target pairs after excluding self-interactions. Cosine similarity was calculated for all candidate pairs. SCENIC was evaluated using the corresponding GRNBoost2 importance scores (Moerman et al., 2019) restricted to the same pairs, without refitting the model. Reference edges were defined as the union of TRRUST interactions and DoRothEA human regulons at confidence levels A–C, with duplicates removed (Han et al., 2018; Garcia-Alonso et al., 2019). The union contained 438 reference edges among the candidate pairs; all remaining pairs were treated as unannotated negatives. Primary performance was measured by average precision (AP), with AUROC as a secondary measure. AP was also normalized by the prevalence of reference edges (438/163,280 = 0.002683). TRRUST and DoRothEA were evaluated separately as sensitivity analyses. Confidence intervals and paired comparisons with GRNBoost2 were obtained from 2,000 TF-cluster bootstrap replicates. These analyses assess recovery of curated regulatory interactions rather than the biological validity of all predicted edges.

##### Motif-supported networks and hub annotation

A separate network analysis used 2,000 matched highly variable genes (Stuart et al., 2019) and a shared human TF list. For each TF, the top 200 non-self targets were selected by embedding cosine similarity or matched-input GRNBoost2 importance and subsequently filtered using pySCENIC cisTarget with hg38 motif-ranking databases and v10nr HGNC annotations (Van de Sande et al., 2020). Motif-supported networks were evaluated for reference-edge support, edge overlap with matched-input SCENIC, and the composition of their top ten TF hubs. Aging-associated TFs were defined using the union of CellAge (Avelar et al., 2020) and GenAge (Tacutu et al., 2018) genes intersected with the TF list. Enrichment among top hubs was tested using one-sided Fisher’s exact tests with Benjamini–Hochberg correction (Benjamini and Hochberg, 1995). A separate sensitivity analysis applied SCENIC to all expressed genes without Top-200 truncation and evaluated the resulting network in the corresponding all-gene universe (Supplementary Fig. S6; Supplementary Table S12).

### Statistical analysis

Cell-level analyses used individual cells, donor-level age analyses used donors, and bulk RNA-seq analyses used samples as the units of evaluation. For cell-type-specific age analyses, predictions were averaged within each donor, cell type and condition. Metrics were not estimated for groups with fewer than three donors.

Donor-level age metrics used 1,000 bootstrap replicates (Efron, 1979; seed 123), each sampling min[N, max(3, floor(0.8N))] donors with replacement. Reported estimates were bootstrap means, with percentile 95% intervals. This procedure resampled approximately 80% of donors per replicate, rather than N donors. Evaluation labels were excluded from downstream predictor fitting, checkpoint selection and threshold selection; cohort-specific scVI fitting was the transductive exception at the representation stage. PERMANOVA used 999 label permutations (Anderson, 2001). GRN enrichment used Benjamini–Hochberg correction (Benjamini and Hochberg, 1995); regulatory-edge AP used the TF-cluster bootstrap described above.

### Cross-task ranking and model-characteristic analysis

The primary endpoints were mean donor-level PCC across two healthy cohorts for T1, mean signed age–pseudotime PCC across six cell types for T2, the fraction of positive disease-associated ΔAgeGap estimates across three disease contexts for T3, mean F1 across four evaluation settings for T4, and pre-motif union-reference AUPRC for T5. Higher values indicated stronger performance for T1, T2, T4 and T5, whereas T3 summarized the directional consistency of disease-associated age shifts.

Within-task rankings used an aggregate-then-rank approach (Wiesenfarth et al., 2021): primary endpoint values were averaged across contexts with equal weights, and methods were ranked from highest to lowest. Ties received average ranks, which were normalized as (N − rank)/(N − 1), where N was the number of methods with complete task coverage (14, 13, 14, 15 and 11 for T1–T5; Supplementary Table S15). Missing results were not imputed. For the cross-task summary, ranks were recalculated among the ten scFMs with results for all five tasks, normalized with N = 10, and averaged with equal weights (20% per task; Supplementary Table S16). Ranking within individual contexts before averaging was retained as a sensitivity analysis. All rankings were descriptive and based on point estimates.

Model architecture, parameter count, context length, pretraining-corpus size, expression encoding and biological priors were compiled from primary publications and official resources (Supplementary Table S13). These characteristics were compared descriptively with benchmark results; individual design features were not evaluated independently.

## Author Approval

All authors have read and approved the manuscript and agree to its submission. This manuscript has not been accepted for publication or published elsewhere.

## Acknowledgments

Shiwen Ni is supported by the Shenzhen Science and Technology Program (JCYJ20250604182917023), Shenzhen Major Science and Technology Project (KCXFZ20240903094007010), GuangDong Basic and Applied Basic Research Foundation (2026A1515060001, 2024A1515012003 and 2023A1515110718).

Hui Li and Rongrong Ji are supported by the New Generation Artificial Intelligence-National Science and Technology Major Project (No. 2025ZD0122702).

---

**Supplementary Figure S1. Cell-type composition of the benchmark datasets.** (A) Composition of six major cell types across the datasets used in Tasks 1, 2, 3, and 5. (B) Cell-type composition of the Task 4 source reference collection and independent OOD datasets. Blue and orange indicate normal (0) and senescent (1) cells, respectively. The cross-domain bulk RNA dataset is excluded because cell-type annotations are unavailable.

**Supplementary Figure S2. Cell-type-specific donor-level age prediction in held-out healthy ZEBRA donors.** Scatterplots show mean predicted versus chronological age for (A) excitatory neurons, (B) inhibitory neurons, (C) astrocytes, (D) oligodendrocyte precursor cells, (E) oligodendrocytes and (F) microglia.

**Supplementary Figure S3. Age–pseudotime concordance across cell types and models.** Heatmaps show (A) Pearson correlation, (B) Spearman correlation and (C) linear-model R² for six cell types and 13 representations. Values are taken from Supplementary Table S7.

**Supplementary Figure S4. Cell-type-specific disease-associated shifts in transcriptomic age prediction.** (A–C) Alzheimer’s disease, multiple sclerosis and schizophrenia, respectively. Within each disease block, rows show PCC, ΔMAE and ΔAgeGap across six cell types. Differences are calculated relative to dataset-matched controls. Error bars indicate 95% confidence intervals.

**Supplementary Figure S5. Additional performance metrics for senescent-cell identification across OOD settings.**

**Supplementary Figure S6. All-expressed-gene native SCENIC sensitivity analysis.** (A) Network sizes for matched-input and all-expressed-gene native SCENIC before and after motif pruning. (B) Fractions of native SCENIC edges supported by the union of TRRUST and DoRothEA A–C regulons. (C) Top 10 native SCENIC TF hubs; red bars identify the TFs ETS2 and RUNX1. (D) Native and matched-input SCENIC shared 607 motif-supported edges, corresponding to 40.1% recall and 2.51% Jaccard similarity. SCENIC concordance indicates methodological agreement rather than regulatory accuracy.

**Supplementary Figure S7. Regulatory-edge recovery assessed by (A) AUPRC and (B) AUROC.** Methods are ordered by AUPRC; fold annotations show AUPRC divided by reference-positive prevalence.

**Supplementary Table S1. Summary of datasets used in the aging single-cell foundation model benchmark.**

**Supplementary Table S2. Models and methods included in the benchmark.**

**Supplementary Table S3. Overall cell-level age prediction performance across models.**

**Supplementary Table S4. Overall donor-level age prediction performance across models.**

**Supplementary Table S5. Cell-type-specific cell-level age prediction performance across models.**

**Supplementary Table S6. Cell-type-specific donor-level age prediction performance across models.**

**Supplementary Table S7. Complete age–pseudotime benchmark metrics.**

**Supplementary Table S8. Overall performance of single-cell foundation models for age perturbation prediction across datasets.**

**Supplementary Table S9. Cell-type-specific performance of single-cell foundation models for age perturbation prediction across datasets.**

**Supplementary Table S10. Senescent-cell identification performance across OOD settings.**

**Supplementary Table S11. Quantitative benchmark of matched-input motif-supported gene regulatory networks across single-cell foundation models and SCENIC.**

**Supplementary Table S12. All-expressed-gene native SCENIC sensitivity analysis and comparison with matched-input SCENIC.**

**Supplementary Table S13. Foundation model architecture and pretraining characteristics**

**Supplementary Table S14. Benchmark cell-embedding and gene-embedding extraction configurations**

**Supplementary Table S15. Context-averaged primary metrics and task rankings.**

**Supplementary Table S16. Five-task composite ranking of ten foundation models.**

**Supplementary Table S17. GRN edge-ranking performance assessed by AUPRC and AUROC. The accompanying workbook includes separate-reference and motif-pipeline sensitivity analyses.**

**Supplementary Table S18. Disease-associated age-shift direction and confidence-interval support.**

