## Supplementary figures and images for "Benchmarking single-cell foundation models for aging biology"

### Supplementary Figure S1

#### Supplementary Figure 1 | Cell-type composition

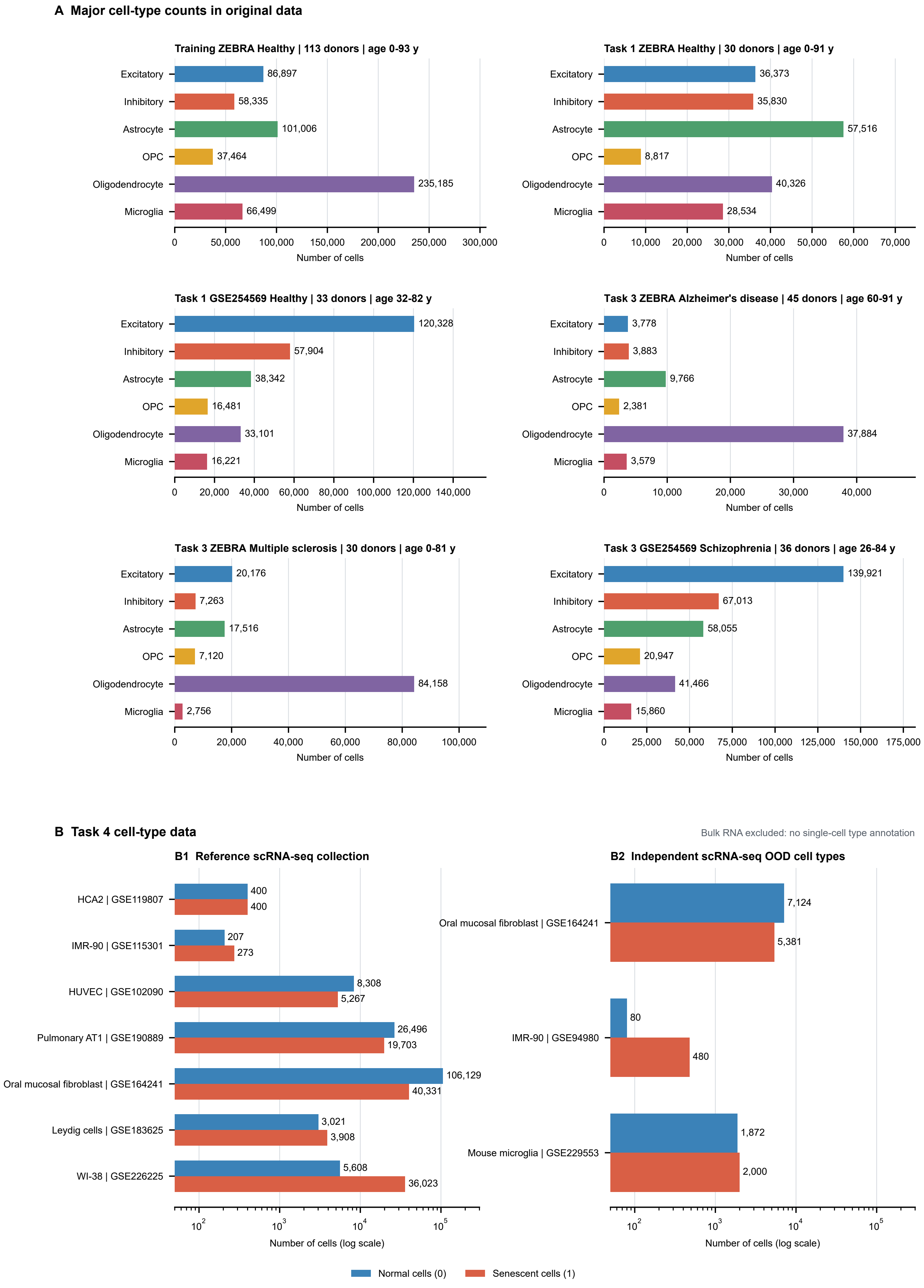

### Supplementary Figure S2

#### ZEBRA | Age prediction

OOD - Unseen healthy donors | Donor level

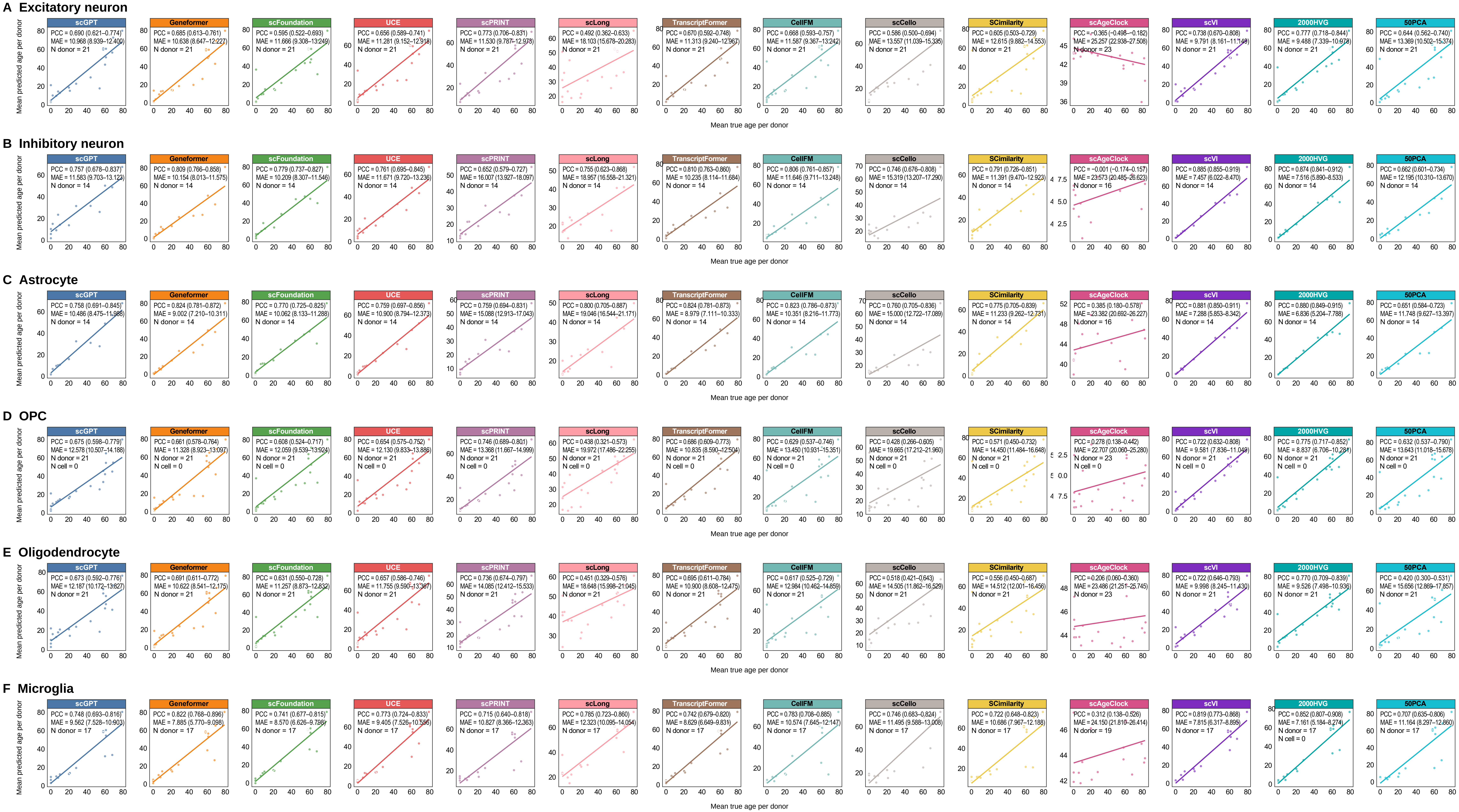

### Supplementary Figure S3

##### Age–pseudotime concordance across cell types and representations

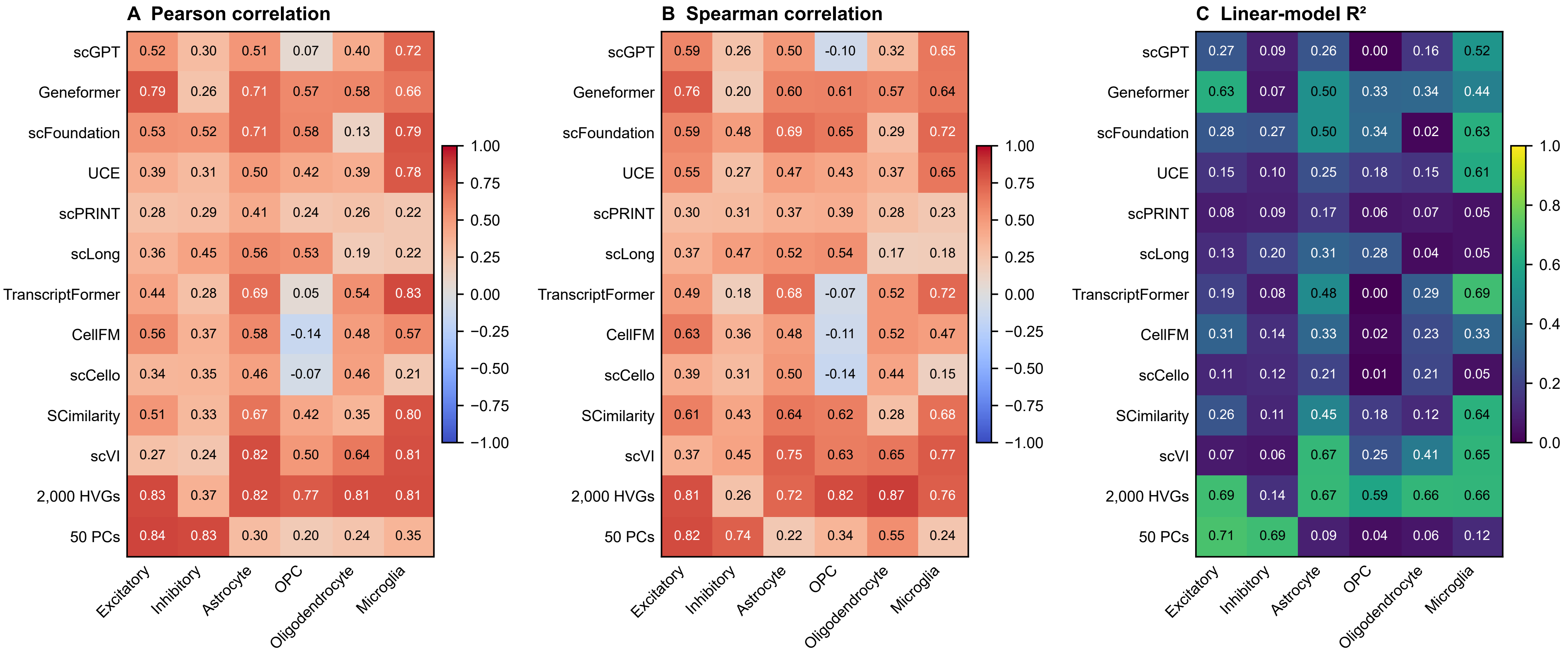

### Supplementary Figure S4

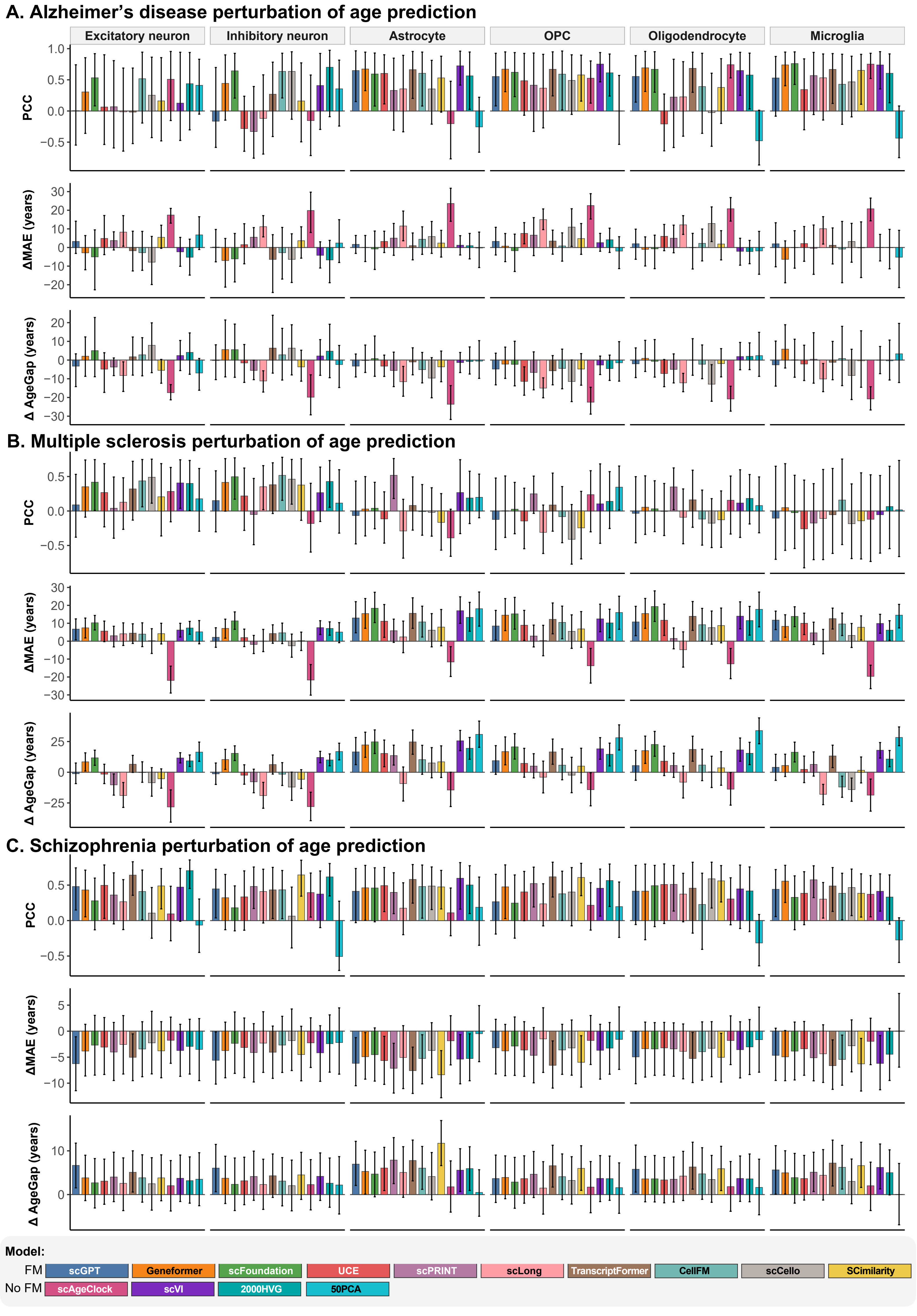

### Supplementary Figure S5

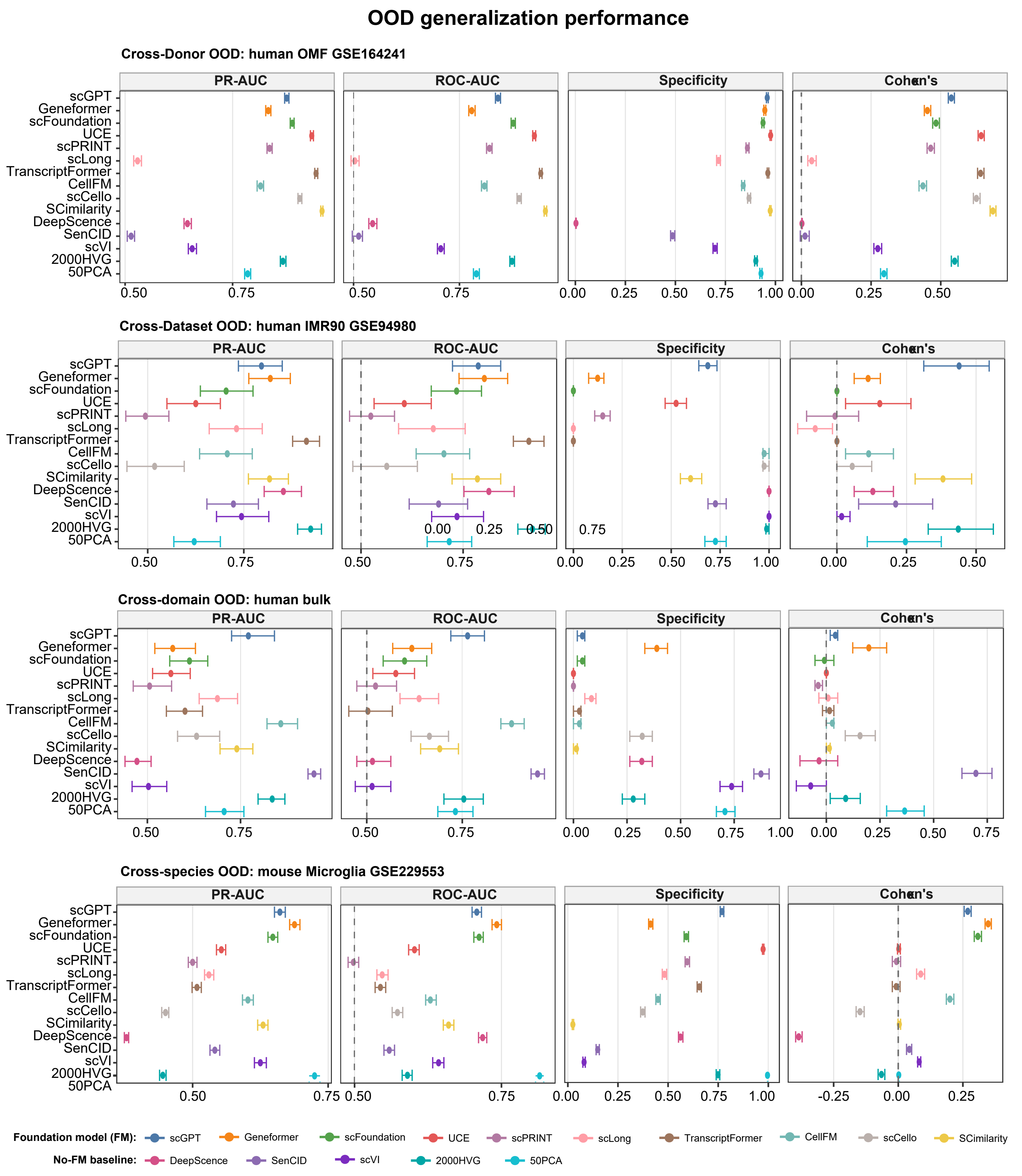

### Supplementary Figure S6

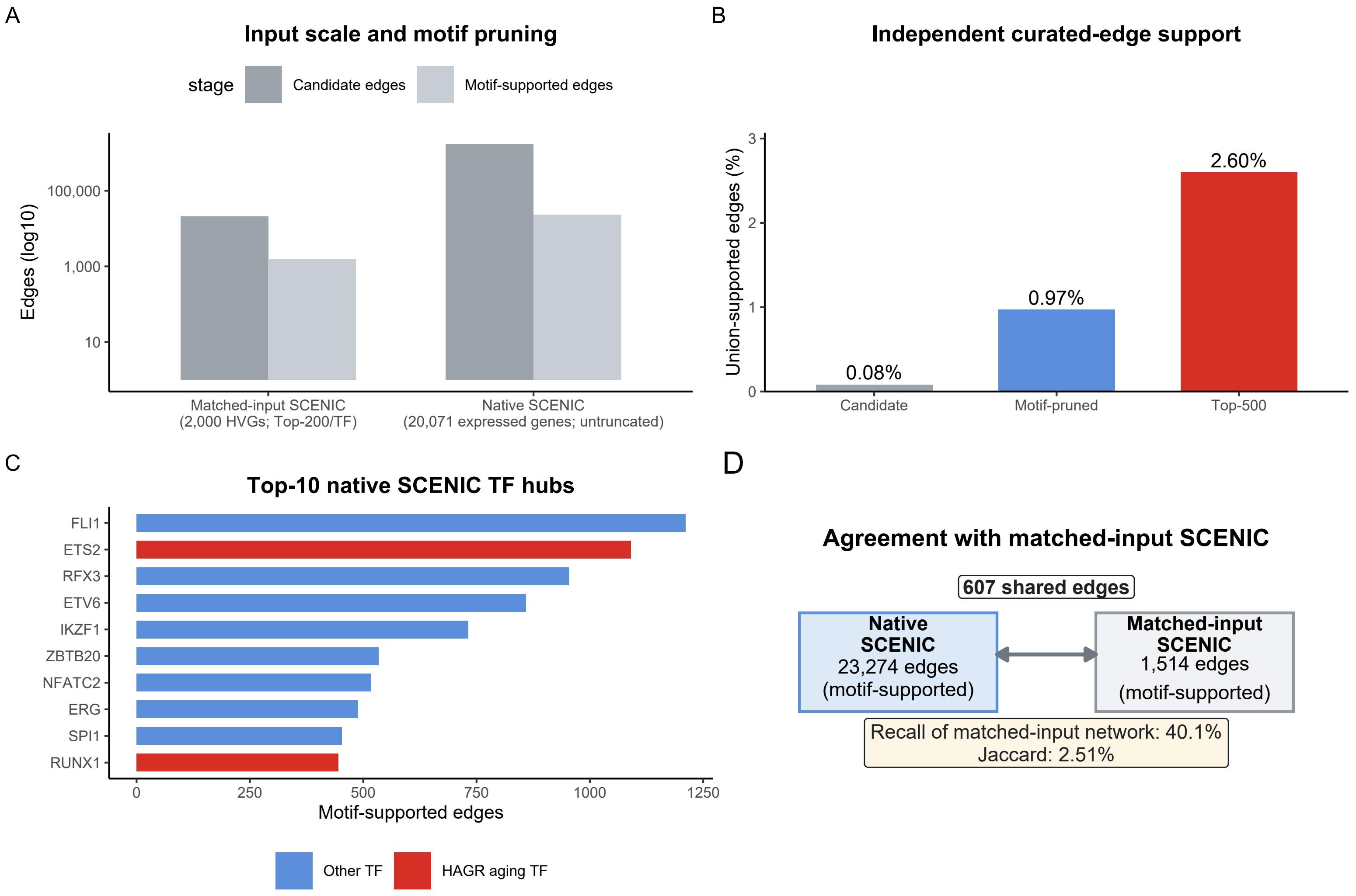

### Supplementary Figure S7

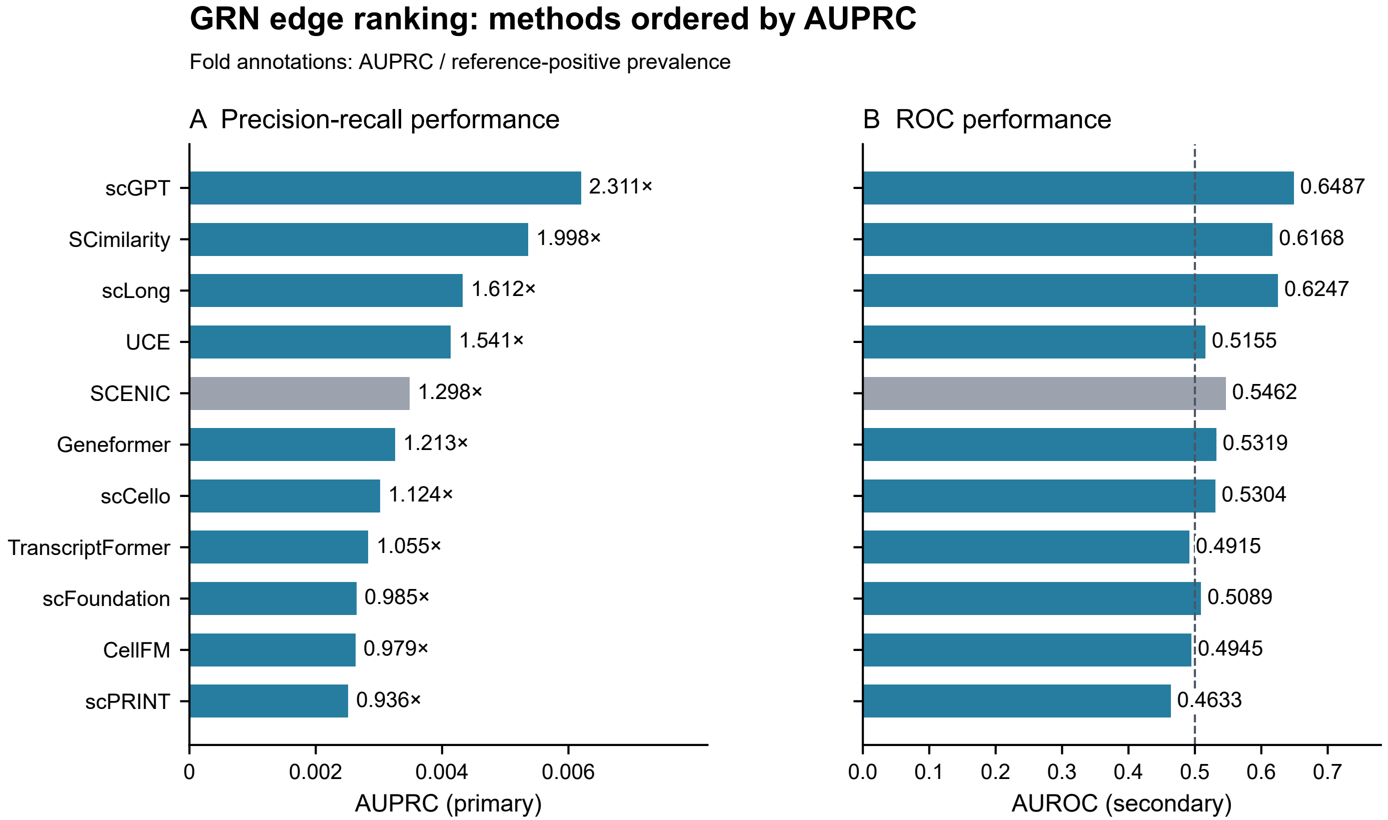
